# Periodic ethanol supply as a path towards unlimited lifespan of *C.elegans* dauer larvae

**DOI:** 10.1101/2021.11.17.468960

**Authors:** Xingyu Zhang, Sider Penkov, Teymuras V. Kurzchalia, Vasily Zaburdeav

## Abstract

The dauer larva is a specialized stage of development optimized for survival under harsh conditions that has been used as a model for stress resistance, metabolic adaptations, and longevity. Recent findings suggest that the dauer larva of *C*.*elegans* may utilize external ethanol as an energy source to extend their lifespan. It was shown that while ethanol may serve as an effectively infinite source of energy, some toxic compounds accumulating as byproducts of its metabolism may lead to the damage of mitochondria and thus limit the lifespan of larvae. A minimal mathematical model was proposed to explain the connection between the lifespan of dauer larva and its ethanol metabolism. To explore theoretically if it is possible to extend even further the lifespan of dauer larvae, we incorporated two natural mechanisms describing the recovery of damaged mitochondria and elimination of toxic compounds, which were previously omitted in the model. Numerical simulations of the revised model suggest that while the ethanol concentration is constant, the lifespan still stays limited. However, if ethanol is supplied periodically, with a suitable frequency and amplitude, the dauer could survive as long as we observe the system. Analytical methods further help to explain how the feeding frequency and amplitude affect the lifespan extension. Based on comparison of the model with experimental data for fixed ethanol concentration, we propose the range of feeding protocols that could lead to even longer dauer survival and can be tested experimentally.

## Introduction

*C.elegans* is a well known free living nematode studied as model organism to address a board range of biomedical questions from genetics, cell biology, and human disease conditions to nematode control [1, 2]. In the context of how organisms may adapt to stressful environmental conditions, *C.elegans* larval stage called “dauer” is of particular interest [3–5]. A developing *C.elegans* larva at L1 stage can turn into an alternative dauer larva developmental stage under harsh environments such as lack of food or high population density [3–5]. To be able to survive these environments, *C.elegans* dauer develop a strong cuticle that covers its whole body, such that most of matter exchange across its body boundary shuts down. As a result, it was long believed that *C.elegans* dauer survive solely on stored lipids and are not able to uptake any carbon source form their environment [3–5]. However, our recent findings [6] showed that *C.elegans* dauer can utilize ethanol as an external carbon source, see Fig 1. Remarkably, at optimal concentrations, ethanol could expand lifespan of dauer larvae two fold for a wild type and up to four fold for some mutants. Ethanol can penetrate across the cuticle, and thus gets channelled in the metabolic pathways of *C.elegans* dauer larvae. The enzymes responsible for the first metabolic steps are SODH-1 and ALH-1, that transform ethanol to acetate which can be activated into acetyl-COA and enters the major metabolic pathways of TCA cycle, glyoxylate shunt gluconeogenesis and and lipid metabolism, thus augmenting the metabolic pathways that dauers use for energy production [5–9]. SODH-1 and ALH-1 are found to be up-regulated in the presence of ethanol, whereas in *sodh-1* mutant, the ethanol is no longer incorporated and does not affect the lifespan of dauer. Experiments with radioactively-labelled ethanol have shown that it can be utilized for the production and accumulation of stored lipids, thus providing an effectively unlimited source of energy to dauer larvae in case of permanent ethanol supply [6].

**Fig 1.**
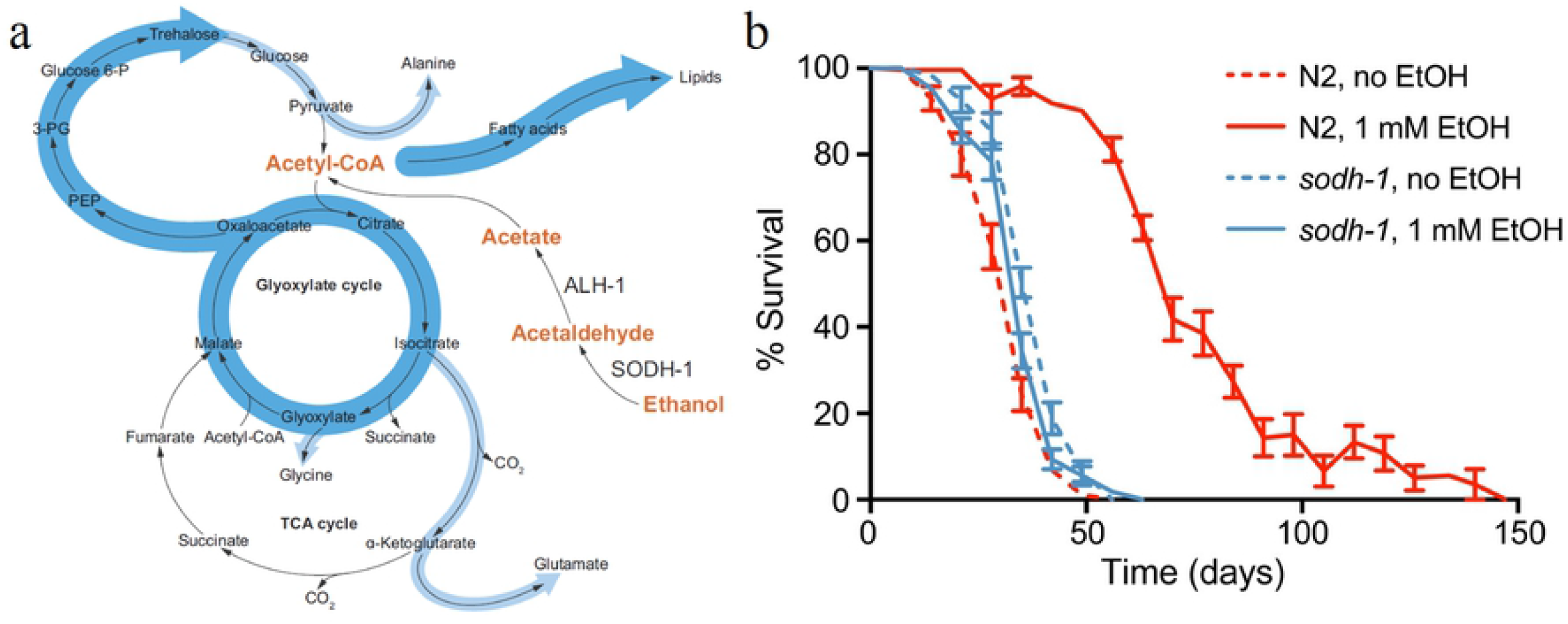
Schematics of the metabolic pathway of dauer larvae in the presence of external ethanol (a), and experimentally recorded lifespans for wild-type and *sodh-1* mutant under ethanol supply and controls without ethanol (b). Figure and data were reproduced from [6] with permission of authors and according to CC BY license.

This led us to the question of why even in the presence of this energy source dauers do exhibit a longer lifespan but then eventually die. To help to answer this question we proposed a mathematical model describing the relation between the lifespan of *C.eleans* dauer and the supplied ethanol, based on the known metabolic pathways of dauer larvae. We assumed that the dauer dies either due to the lack of energy or due to the accumulation of some, not yet identified toxic compound(s) [6] that could resemble the so-called “lipitoxicity” factors in mammalian systems [10–12]. As experimentally observed, the death of worms was preceded by deterioration of mitochondria. We also assumed that these two mechanisms lead to mitochondria damage and then to death. This model was very successful in explaining experimental data on lifespans of dauer and various mutants in presence and without ethanol.

While identifying the exact toxic component that limits the lifespan of dauer is still an ongoing research project, we were interested to explore whether or not the lifespan could be extended even further. To this end, we assume there are two self-recovery mechanisms, namely regeneration of mitochondria and detoxification, and test what they lead to. These two mechanisms alone still result in dauer’s death if feeding protocol is constant. However, when we use periodic supply of ethanol in the model, an unlimited lifespan can emerge according to the numerical simulation. By comparing model predictions with existing data, we also suggest feeding protocols that now can be directly tested in future experiments on dauer.

## Mathematical model

A simple model of the metabolic network of *C.elegans* dauer larvae was introduced in [6] and accurately reproduced the lifespans of dauer with and without ethanol for wild type worms as well as for various mutations. The framework of the model follows the largely coarse-grained metabolic pathway of dauer. All the chemical components falling into the category of “available energy” are combined and called “acetate”, which is the central representative component of this category. Similarly, the components corresponding to “stored energy” and “consumed energy” are denoted as lipids and carbohydrates respectively. Acetate and lipids could transform between each other as the balance between free and stored energy. At the same time, acetate continuously transforms into carbohydrate unidirectionally to support the main functions of an organism including mitochondria. If production of carbohydrates drops below a certain minimal threshold, mitochondria start to get damaged and the dauer dies. In presence of ethanol, acetate gains an influx proportional to its concentration. During the process of releasing stored lipid, toxic compounds are produced as a side product, as the second major reason damaging the mitochondria alongside the lack of carbohydrate production.

Our model also included the effect of genes that were identified as regulatory factors through genetic experiments with loss-of- or reduction-of-function mutations [6]. *daf-2(e1370)* is the mutant that always enters dauer state without the need of stress conditions. We will refer to it as a control and use it as a proxy for the wild-type dauer under stress, since the experimental results show very little difference between *daf-2* and wild-type dauers. Loss-of-function mutation in the *aak-2* /AMPKα in *daf-2(e1370);aak-2(gt33)* double mutants causes an enhanced lypolisis rate, which leads to a reduced lifespan as compared to control strain under both feeding conditions with and without ethanol [7, 13]. A reduction-of-function allele in the class I PI3-kinase *age-1(hx546)*, on the other hand, is supposed to have the reduced lipid synthesis rate. This assumption is based on experimental results with the mutant *age-1* in dauer state [6]. Its lifespan is similar to control dauer when there is no ethanol supply, but has a large increase when the ethanol is supplied [6].

The goal of this work is to identify potential ways of how the dauer could survive even for a longer time. Thus here we consider mechanisms by including which the model will be able to produce an unlimited lifespan while still remaining consistent with the results of the previous experiments. There are two essential and rather natural mechanisms that have been omitted in the original model [6] while having a potential for lifespan extension: detoxification [15, 16] and a possibility for mitochondria to regenerate [17–19] (see green arrows in Fig.2). We will show in the following, that the model containing these two mechanisms predicts the possibility for lifespan extension under periodic supply protocol of the ethanol.

**Fig 2.**
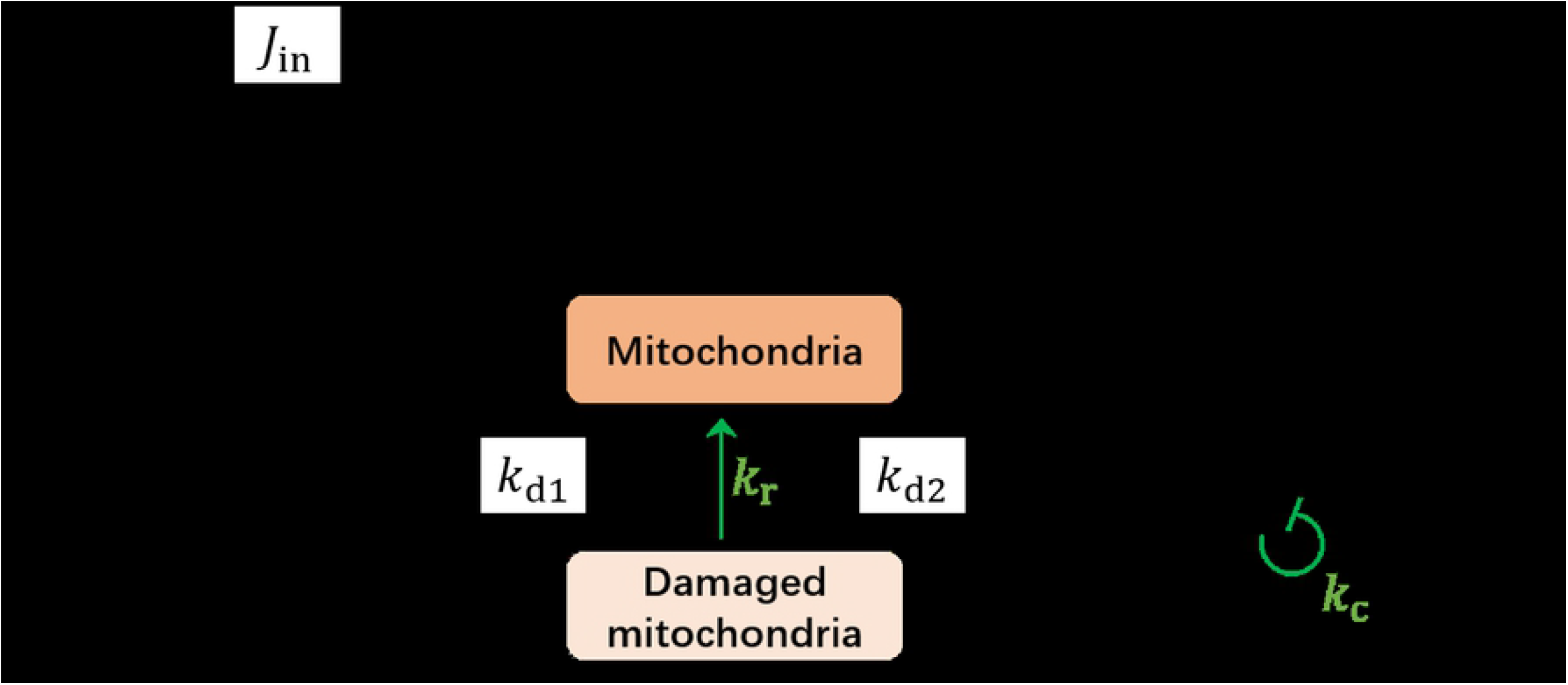
Schematics of the mathematical model of the metabolic network of *C.elegans* dauer larvae. Externally supplied ethanol is transformed into the acetate with the rate *j*_in_, which can be either constant or varies in time depending on the supplied ethanol concentration. Acetate (ch. acetyl-CoA in Fig 1) is then used either for energy production and carbohydrate synthesis or can be stored in lipids. Mitochondria takes damage from lack of carbohydrate production or accumulation of toxic compounds as the product of lipolysis.

To demostrate this, we first formalize the schematics in Fig.2 into the system of ordinary differential equations that describe the chemical reaction network of ethanol metabolism [14]:

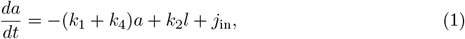

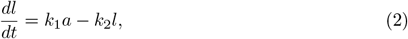

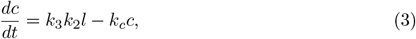

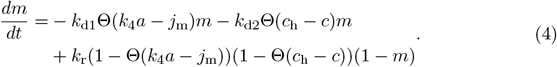

Here “*a*” and “*l*” denote the concentrations of the acetate and lipids respectively, while “*c*” represents the concentration of toxic compound(s), and *m* designates the well-being of mitochondria. The consumption of acetate for simplicity is assumed to be unidirectional (not explicitly modelled in the system) with the rate *k*_4_, but acetate can also be stored in lipids, see Eq.(1). In presence of external ethanol, an influx *j*_in_ of acetate is included as the source. This influx *j*_in_ is assumed to be proportional to the external ethanol concentration. Lipids get created from acetate with rate *k*_1_ while they are released through lipolysis process with rate *k*_2_, Eq.(2). The toxic compound(s) *c* is produced as the side product of lipolysis with proportionality factor *k*_3_ and spontaneous degradation rate *k*_c_, Eq(3). The variable *m* ranges from 1 to 0 denote the well-being of mitochondria, where *m* = 1 means a fully functional mitochondria and *m* = 0 means a fully damaged one. Mitochondria can be damaged with a rate *k*_d1_ if the carbohydrate production *k*_4_*a* falls below the minimal required “energy” flux *j*_m_, or with a rate *k*_d2_ when the toxic compound *c* accumulates above a certain threshold concentration *c*_h_ (Θ in the equation is the Heaviside step function). There are many known mechanisms of mitochondria surveillance and maintenance [17–19]. Here for simplicity, we suggest a phenomenological law of mitochondria recovery, where the mitochondria regenerates its current damage level (1 *− m*), provided it is not suffering from any further damage with a constant regeneration rate *k*_r_.

While most of reaction rates in the above equations are considered constant for simplicity, some rates do depend on variables. First example is the linear dependence of *k*_4_ on *m*, which assumes that the energy production requires functional mitochondria:

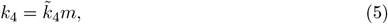

where 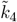 is a constant. Another non-constant rate is *k*_1_ quantifying acetate-to-lipids conversion. It reflects the fact that each dauer has a storage limit capacity *l*_s_ (it cannot accumulate unlimited amounts of lipids):

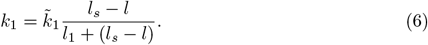

Here *l*_1_ is a characteristic lipid concentration at which the conversion starts to saturate and 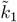 is a constant. Finally, we also assume that *k*_2_ has the functional form of Michaelis-Menten reaction [14]

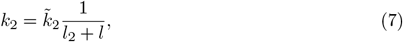

where 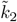 and *l*_2_ are constants. If in the above equations, we set *k*_*r*_ = 0 and *k*_*c*_ = 0 we will recover the system studied in [6].

The model including self-recovery mechanism, however, should also reproduce lifespan of dauers with and without ethanol, as well as different mutants as was observed experimentally [6]. This also means that this model should result in finite lifespan under constant ethanol supply. However, the novel possibility for lifespan extension may now emerge for a non-constant feeding, where the supplied ethanol concentration varies in time, for instance, according to a sinusoidal protocol. We next show by using numerical simulations that the model reproduces experimental observations under constant feeding and predicts the lifespan extension under periodic feeding protocol.

## Results

### 1 Constant ethanol supply

We first check if the model with the self-recovery can recapitulate experimental observations with a constant ethanol supply. Parameters used in the simulations were chosen by checking whether the lifespan ratios between mutants (*daf-2, daf-2;aak-2* and *age-1)* with and without ethanol generated by simulations fit the previous experimental results [6]. When one set of parameter is considered as the control strain without ethanol feeding, the corresponding parameter set of *daf-2;aak-2* mutant is defined by increasing the value of 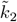 while keeping other parameters unchanged. Similarly, *age-1* mutant has a reduced 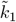 constant. Three strains under two feeding conditions give rise to total of six sets of parameters including the starved control strain as the baseline parameter set. We can call these sets as a parameter collection. A parameter collection of the model is said to reproduce the experimental observation if all the ratios of lifespans produced by its parameter sets reproduced the experimental observed value. The Fig 3 shows the reproduction of experimental observed lifespan ratio by the model with self-recovery.

**Fig 3.**
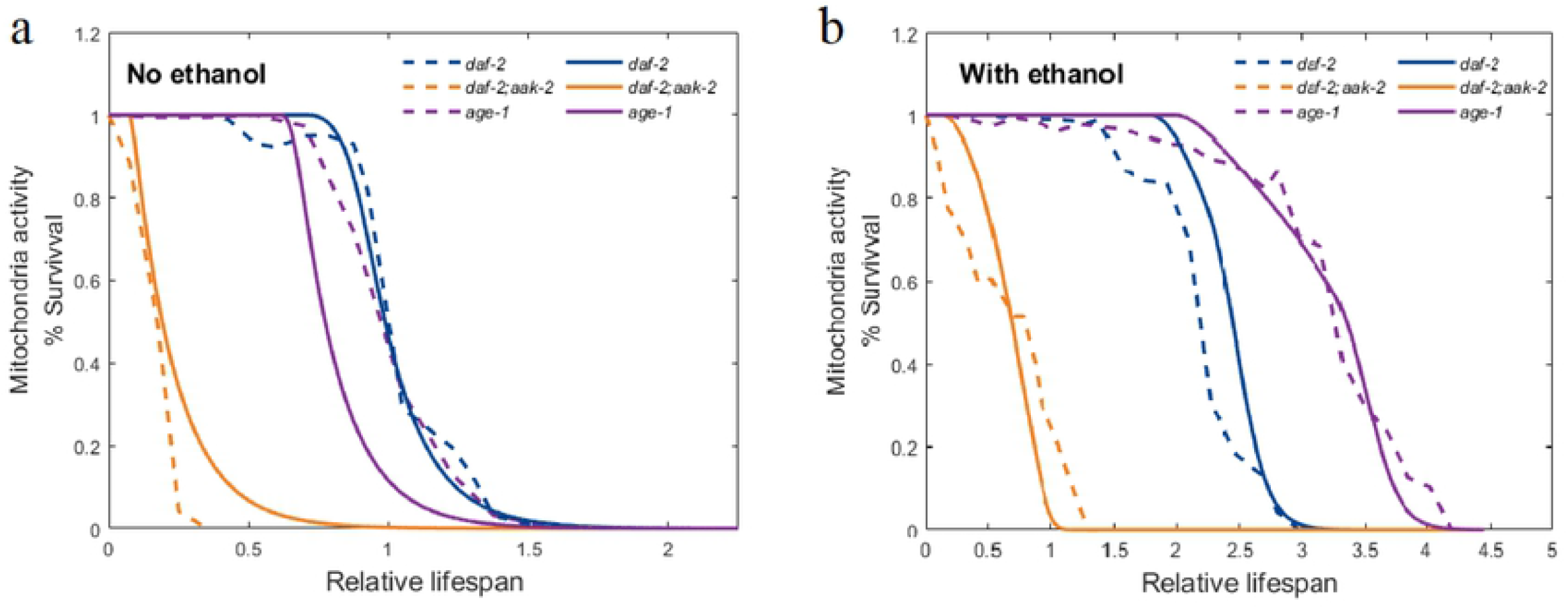
Comparison between experimental observed lifespan of dauer strains (dashed lines) and the corresponding simulation result (solid lines) without (a) and with ethanol feeding (b). The ratio of 1 is set at 50% of control strain.

The detailed dynamics of one of the parameter sets, for control strain with and without feeding is shown as an example in Fig 4. When there is no feeding, dauer brakes down storage lipids to keep its acetate level and thus the carbohydrate production rate. As the lipids run out, the mitochondria is damaged for lack of carbohydrate production and results in the death of dauer. When ethanol is supplied at a sufficient level, starvation becomes no longer a concern. However, the toxic compound continuously accumulates and as it goes beyond the threshold at some point, the mitochondria start to take damage and finally the larvae die. The details of the simulation including the numerical methods [14, 20, 21] and parameters are provided in the Appendix I.

**Fig 4.**
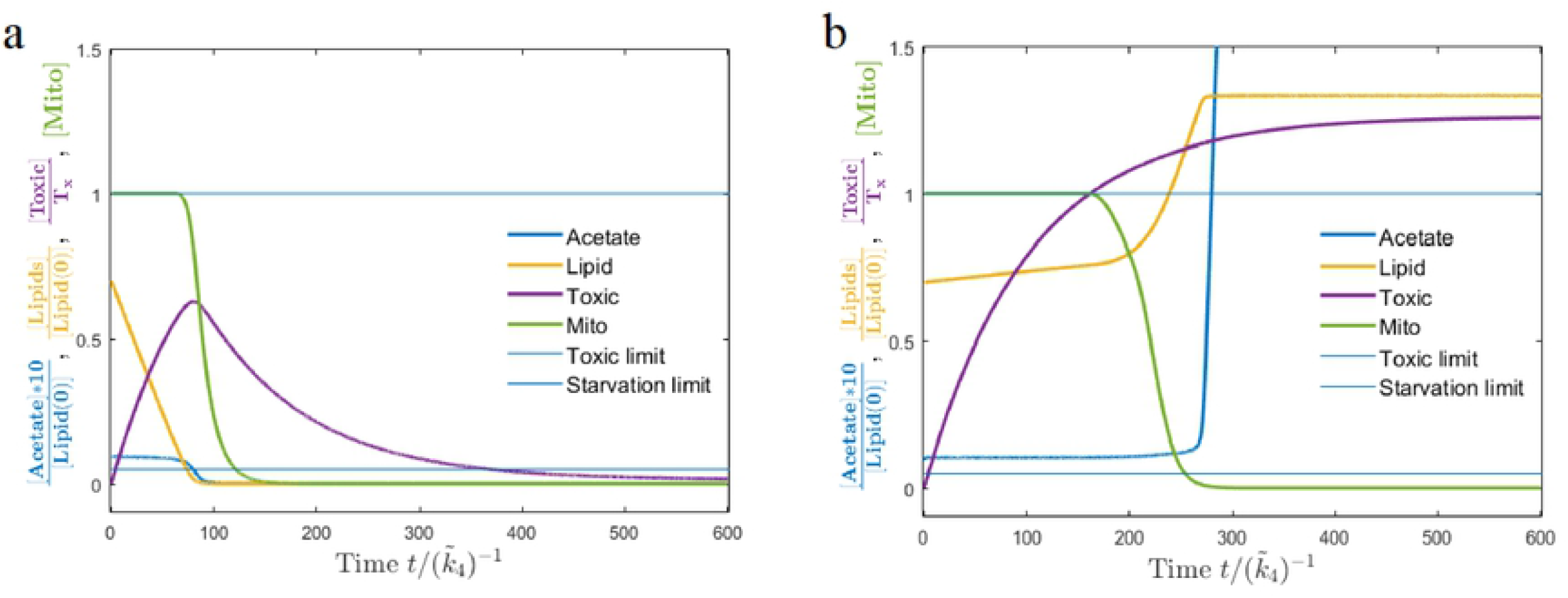
Dynamic of the model corresponding to control strain without (a) and with ethanol (b). Without ethanol supply the dauer consumes the stored lipids and dies due to starvation, while with ethanol the dauer dies due to accumulation of toxic compounds.

As the above two examples show, starvation or accumulation of toxic compounds is the reason for mitochondria damage and the resulting death of larvae. We can demonstrate more generally the condition for the finite lifespan of dauers for a constant ethanol concentration.

Fig 5 shows the lifespan of dauer as a function of the external ethanol influx. If either starvation damage *k*_*d*1_ or toxic damage *k*_*d*2_ is removed, the dauer may have an infinite lifespan when the ethanol concentration is sufficiently high or low, respectively. If, however, these two lifespan vs. influx curves intersect at some value of *j*_in_, lifespan will always remain finite, as for any given ethanol concentration and the corresponding influx, there will be at least one reason that the dauer dies within limited time determined by *k*_*d*1_ or *k*_*d*2_.

**Fig 5.**
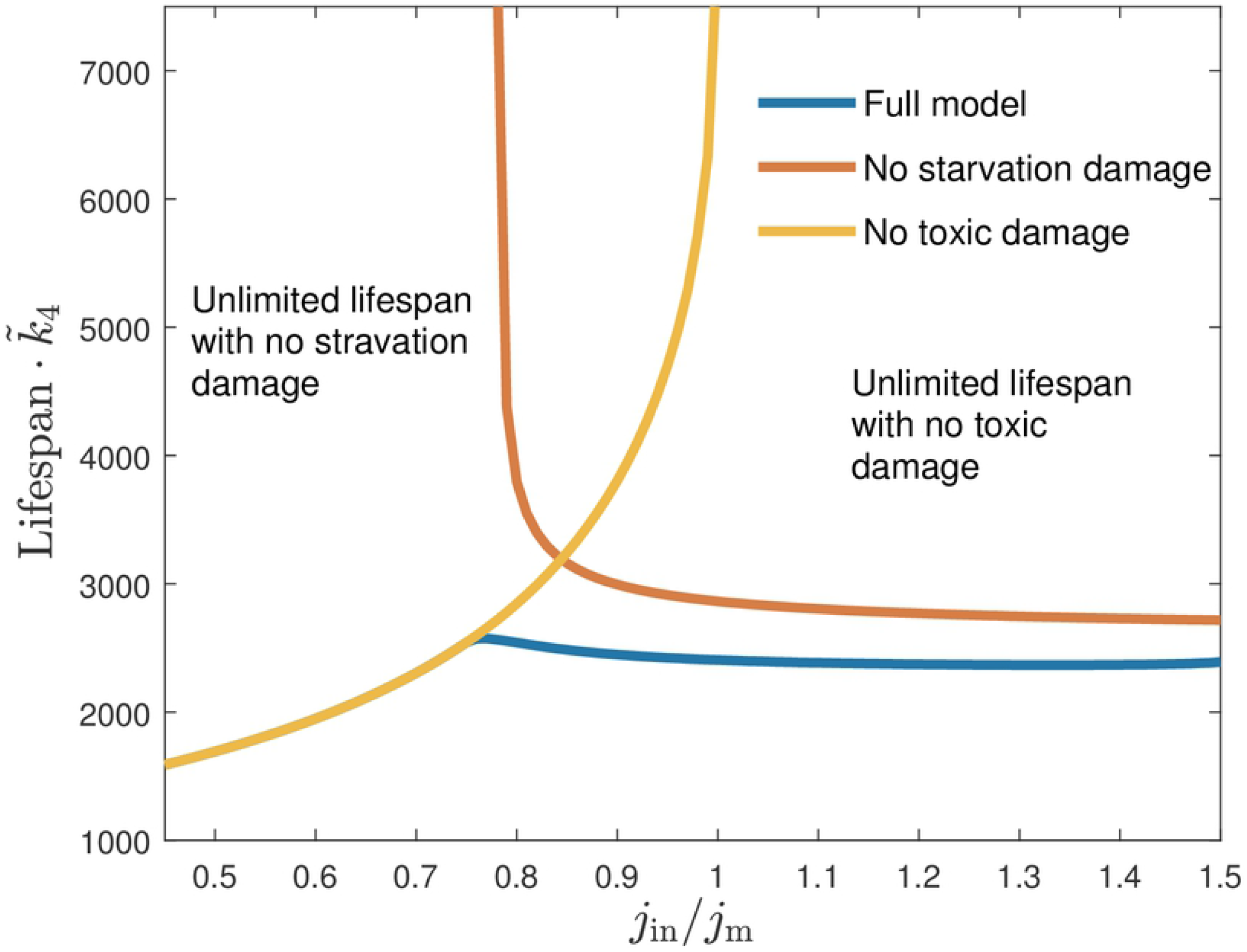
No lifespan extension is possible when larvae are fed with constant ethanol influx. The influx is normalized by minimal energy flux *j*_*m*_ required for mitochondria wellbeing.

If the two curves (with each of the damage removed) do not intersect, and they thus would form boundaries of a domain in between where the value of *j*_*in*_ would support an infinite lifespan. As we mentioned above, in experiments, the dauer survives always the finite time in presence of ethanol, thus defining for us the parameter range that has to be chosen in simulations.

### 2 Periodic ethanol supply

The above results show that ethanol supply keeps mitochondria operational, but the accumulating toxic compounds damage the mitochondria. Here we hypothesize that periodic ethanol supply might be the key to an unlimited lifespan of dauer. While periods of supply might be used to replenish lipid storage and repair mitochondria, the periods of no feeding can be used to degrade the accumulated toxic compounds. We now test this hypothesis numerically. For simplicity, we use a sinusoidal feeding protocol with a feeding amplitude *A*, feeding frequency *ω*_*E*_ and a positive baseline value *j*_0_:

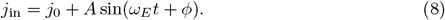

With a proper parameter choice the numerical simulations of the model show that the mitochondria damage and regenerate periodically until the end of simulation, no matter how long these last.

Indeed, this situation becomes possible when parameters are tuned such that the periodic feeding permits the worm to accumulate toxic compounds while intaking ethanol and fuelling mitochondria but then remove them with a diet at the cost of some mitochondria damage, which can, however, be regenerated during the next intake cycle. These simulations suggest that periodic feeding protocol does provide a theoretical possibility of an unlimited lifespan extension (Fig 6). We next investigate in more detail how this effect depends on model parameters.

**Fig 6.**
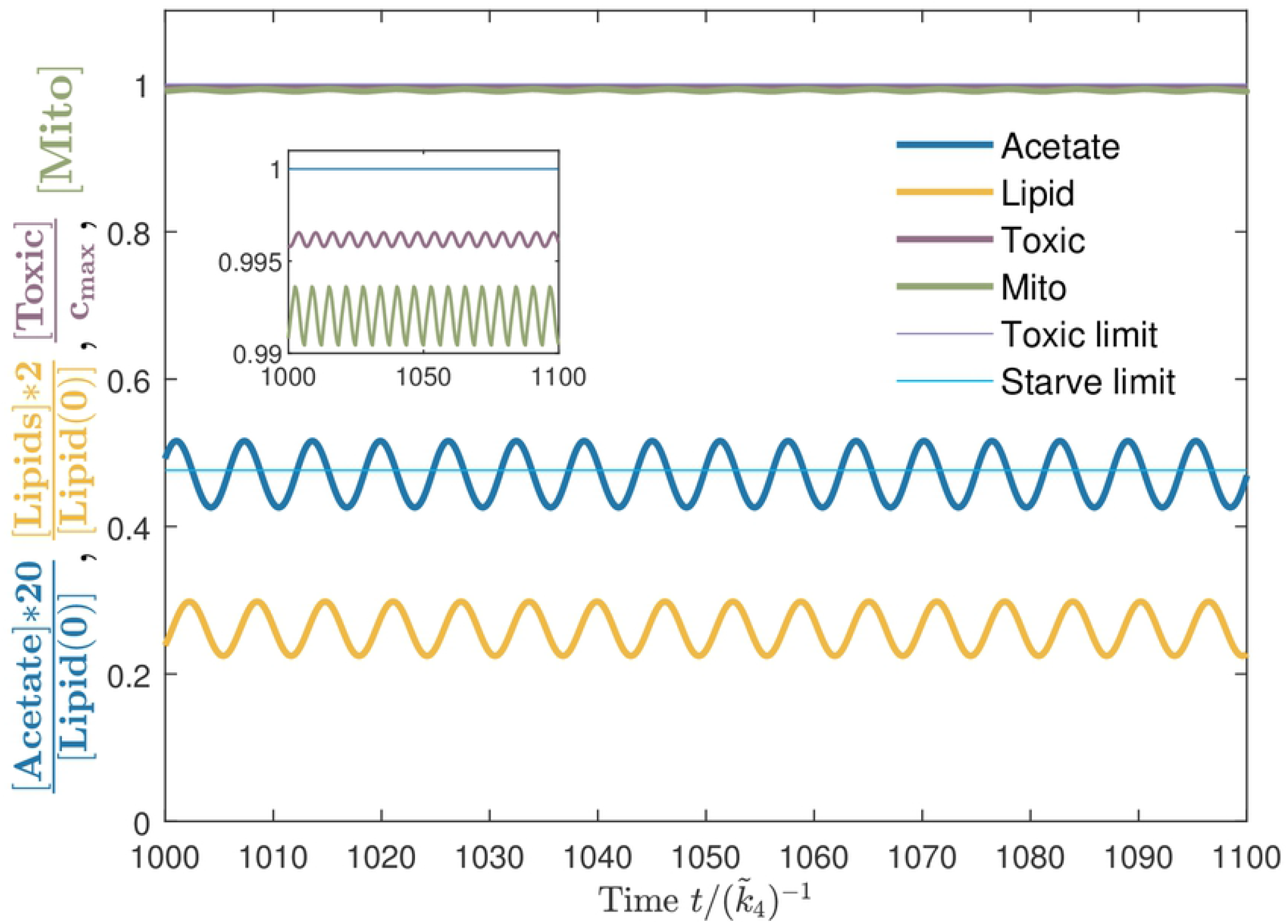
Example of a lifespan extension as a result of periodic feeding. The baseline value *j*_0_*/j*_*m*_ = 0.8, while *A* = *j*_0_. The feeding frequency *ω*_*E*_ is the same as the oscillatory frequency of acetate level *a* shown in blue. The inset shows oscillations of toxicity near but below the toxic limit, while mitochondria are almost normal.

Numerically, an unlimited lifespan can be defined as the survival until the end of simulation regardless of the simulation time. However, in practice the time for which we can observe the system is always limited. Thus we set a certain threshold value *T*_max_ for the survival time. If a dauer survives until *T*_max_ in simulation, we say the lifespan of dauer is unlimited under this set of parameters. Our analytical considerations also suggest that there may exist a true infinite lifespan given certain set of parameters in the model.

### 3 Effect of feeding parameters

A single simulation does not reflect the whole picture, but only indicates a possibility. To quantify the robustness of lifespan extension, we defined a new value “range” *w* as the size of the interval, within which the baseline influx *j*_0_ may vary, so that the dauer exhibits an unlimited lifespan, see Fig 7. The range *w* of this baseline interval thus quantifies the ability of a certain set of parameters (*ω*_*E*_, *A*) to support the extension of lifespan. We note that the lifespan seems to go through a sharp transition from finite to an infinite value. First when approaching from the side of low baseline value, and second when approaching from the side of large baseline levels. The transition at low influx seems to be a jump like switch (as we could only test numerically). The high influx condition is amenable to analytical analysis and we could show that it has a shape of a logarithmic divergence (see Appendix III). By quantifying the survival ability with the value of *w* and studying its dependence on feeding parameters *ω*_*E*_ and *A* would eventually help us to identify the optimal experimental conditions where the lifespan extension of dauer could be tested.

**Fig 7.**
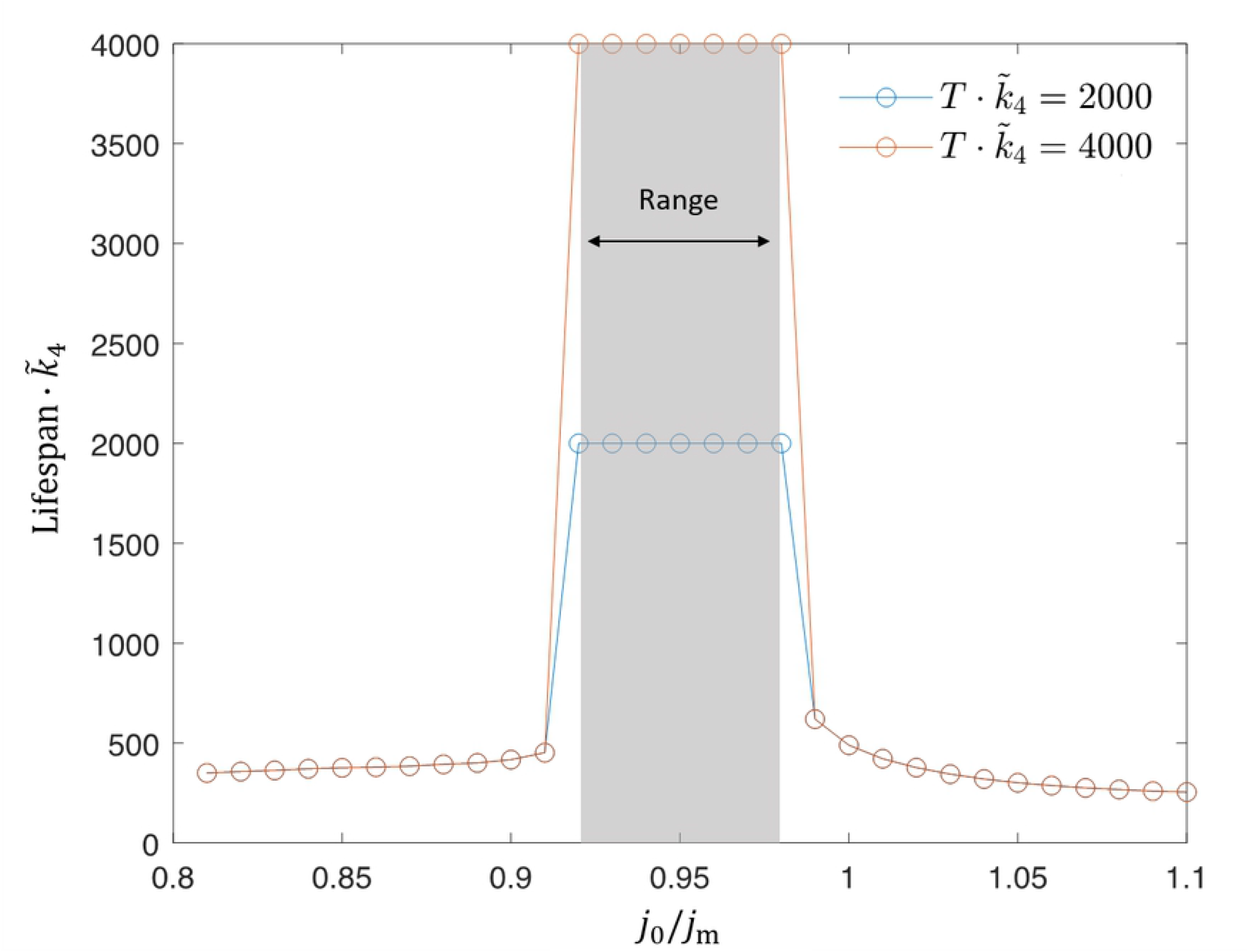
Range *w* is the size of an interval of *j*_0_ where the dauer experiences lifespan extension. Its value is obtained numerically by scanning over the baseline value *j*_0_ while keeping *ω*_*E*_ and *A* fixed. For each *j*_0_ we run multiple simulations corresponding to various initial lipid storage *l*(0), and take the largest *w* from those simulations as the range *w* corresponding to this *j*_0_. The baseline value *j*_0_ can be smaller than the minimal energy flux *j*_m_ as the oscillations with large enough amplitude *A* make *j*_in_ larger than *j*_m_ for a certain fraction of the feeding period. Here we also show that the range does not strongly depend on the chosen threshold, 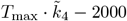, 4000, which is approximately 20 and 40 times greater than the survival time of control strain without ethanol.

We next plot the range *w* as a function of feeding amplitude *A* and frequency *ω*_*E*_, see Fig 8.

**Fig 8.**
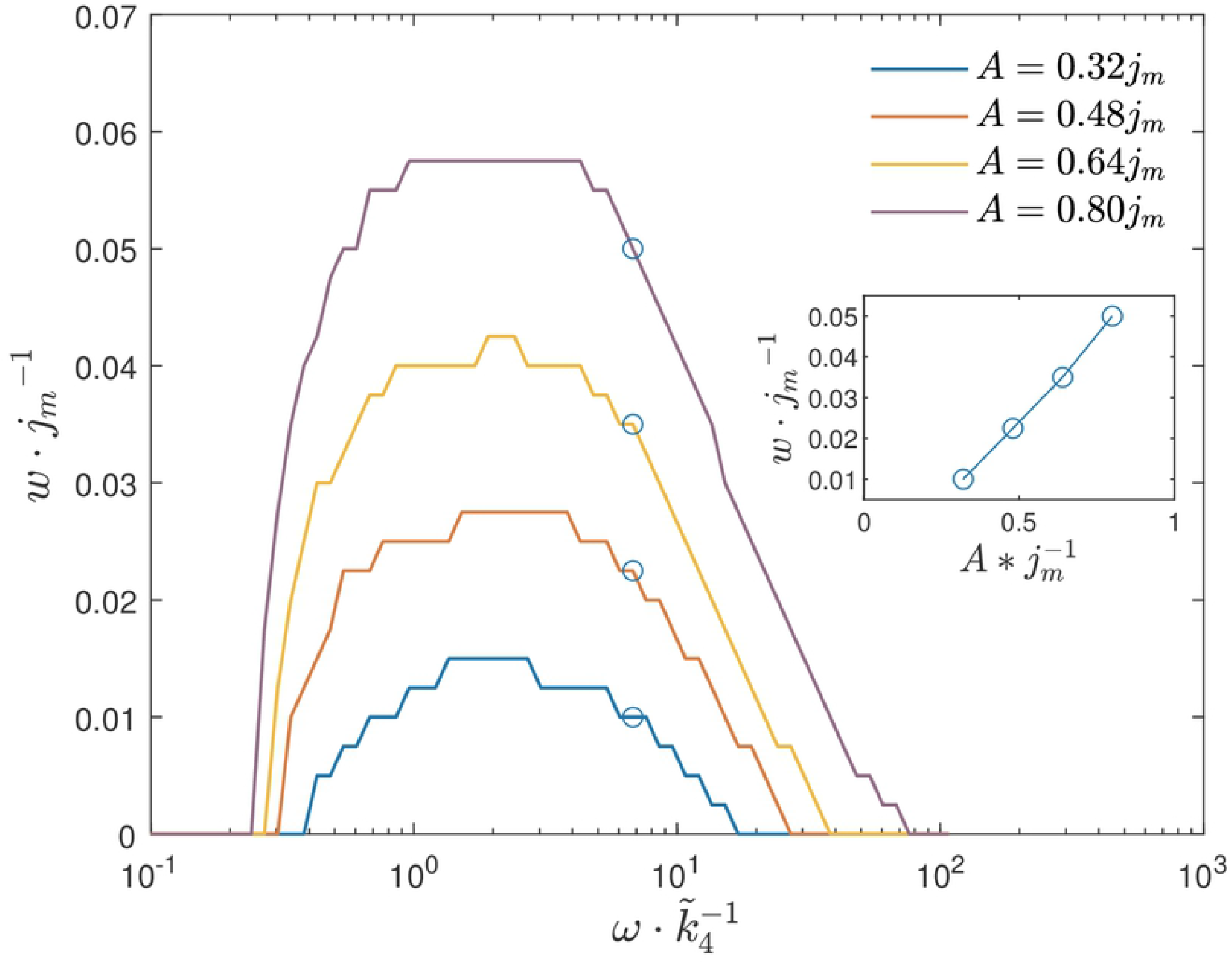
Range *w* as a function of the feeding frequency *ω*_*E*_ for different values of amplitude *A*. The inset shows a linear-like dependence of the range on amplitude in the region of high feeding frequencies.

According to the simulation results, we see that the lifespan extension is possible when the feeding frequency is within a certain interval. The experiment with the feeding frequency *ω*_*E*_ corresponding to the maximal range *w* is expected to give highest chance to observe the effect. The simulations also suggest that the range *w* grows with feeding amplitude *A* in an almost linear way if the feeding frequency is high enough. This is because at high frequency region the range *w* is proportional to the oscillation amplitude of acetate, which can be explained by an approximate analytical solution. (For details of the analysis, see Appendix. II.)

The range-frequency curves can potentially help us to identify suitable feeding frequency and amplitudes for which it is most likely to observe lifespan extension in experiments. To do so, we still need to connect our mostly dimensionless equations to realistic parameters. This is not too straight-forward since not all parameters of the enzymatic kinetics as well as chemical concentrations in the dauer were measured yet. However, for the case of the feeding frequency, we may take a short-cut, where we can determine the timescale by equalling the control lifespan without ethanol in the model defined as time where *m* falls to, for example 0.5 to that in experiment defined as respective 50% survival and restore all reaction rates in real time units. Also the feeding amplitude is simply set as large as possible (see below) so there is no more information needed. Fig 9 shows the range vs period (given in hours) relation for control strain and *daf-2;aak-2* mutants under maximal feeding amplitude. The maximal feeding amplitude is defined as *A* = min[*j*_0_] (i.e. the smallest *j*_0_ among all *j*_0_ used in scanning, such that the influx *j*_in_ is always positive). Another definition *A* = *j*_0_ for all *j*_0_ is also possible and leads to similar results.

**Fig 9.**
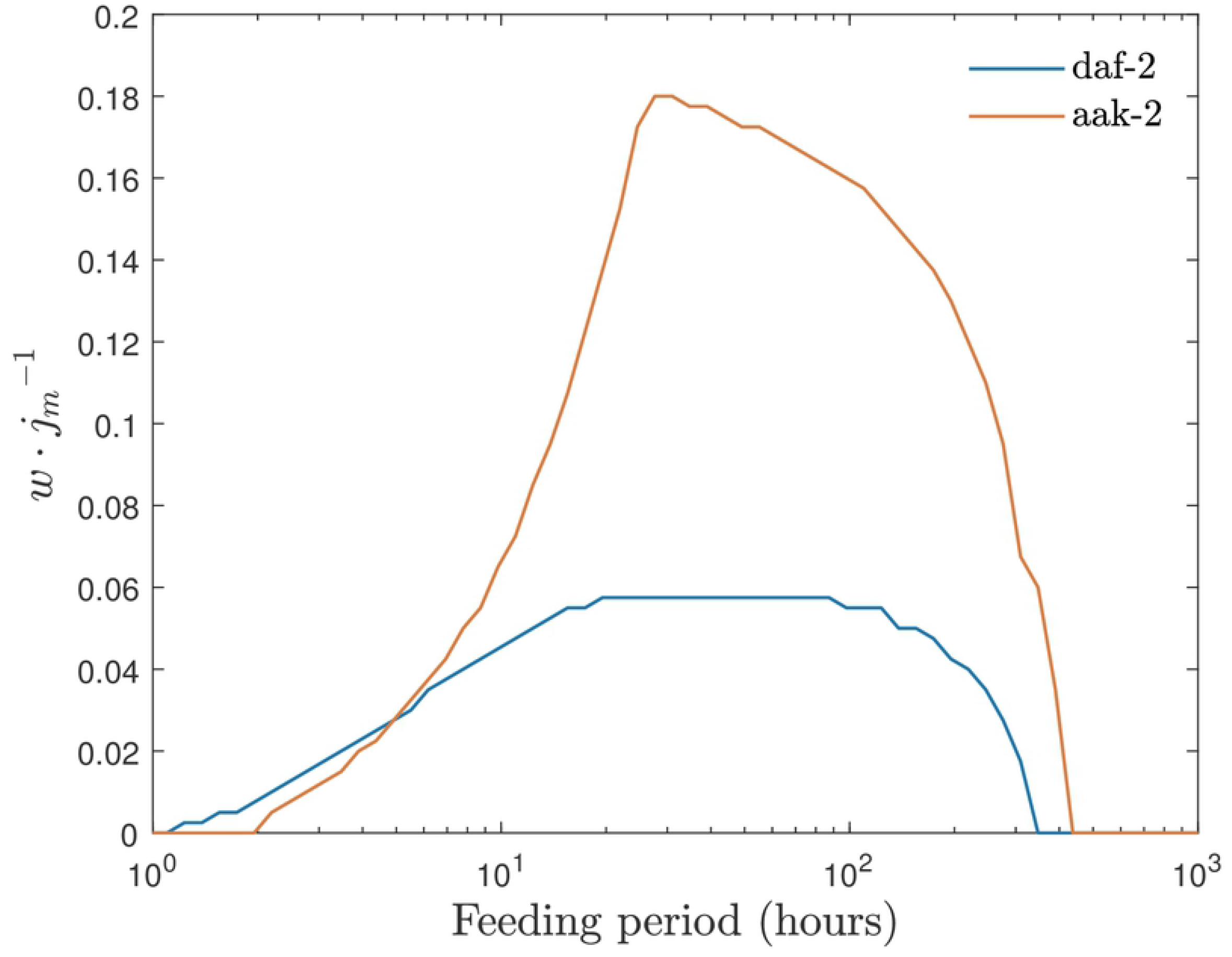
Range of the baseline ethanol supply levels where we expect to see unlimited survival of dauer larvae shown for control and *daf-2;aak-2* strains as a function of the feeding period *T* = 2*π/ω*_*E*_. Here it is assumed that the amplitude *A* takes the value of min[*j*_0_].

These simulations not only suggests an optimal feeding period for both strains, but also indicate that *daf-2;aak-2* mutant is a better option for experiment, for not only the larger range value but also the smaller optimal feeding period. It requires feeding period of the order of 10 hours (so the media for larvae can be changed once a day) and the effect should be seen much earlier, as the original lifespan of *daf-2;aak-2* is much shorter and thus overall a shorter experiment could be carried out. 237

## Discussion & Conclusion

Previously we have shown that the lifespan of *C. elegans* dauer larvae can be greatly extended due to metabolism of externally provided ethanol. With the help of the mathematical model of this metabolic pathway we proposed that the lifespan remains limited due to accumulation of toxic compounds resulting from process of lypoisis. So far, however, we have neglected the possibility of mechanisms that help dauer to recover from this damage. Therefore, two biological self-recovery mechanisms, namely the detoxification and mitochondria regeneration, were introduced into the model. Importantly, despite self-recovery mechanism, for constant ethanol supply, model reproduces the experimental observations of extended but limited lifespan.

However, when feeding protocol is periodic, an unlimited lifespan can emerge. The possibility of the unlimited lifespan can be explained by the switch between two feeding phases, where the first one at high ethanol concentration repairs the mitochondria at the cost of toxic compounds accumulation while the second one, at low ethanol concentration, has the toxic compounds degraded but also damages the mitochondria slightly. For this process to keep the dauer surviving thus requires both, mitochondria regeneration and toxic compounds detoxification mechanisms, to function.

To characterize the unlimited lifespan predicted by the model systematically, we defined a range of baseline feeding fluxes, which quantifies the ability of a certain set of feeding parameters to support the unlimited lifespan. The dependence of this range on feeding frequency and amplitude were studied numerically with some supporting analytical arguments. This dependence combined with previous data helped us to suggest suitable feeding period and amplitude that can now be tested experimentally. If lifespan extension of dauer larvae appears under periodically supplied ethanol, that could be a confirmation of our hypothesis about the existence of recovery mechanisms.

This study treats the identity of the toxic compounds open and does not specify the concrete mechanisms of mitochondria recovery and detoxification. Ultimately for our comprehensive understanding of dauer larvae lifespan extensions mechanisms and generalization of those to other organisms we need to push towards identifying the exact biological players of toxicity and recovery competition.

## Supporting information

**S1 Appendix.I Numerical algorithm and model parameters**. Summarize the numerical methods and parameter used in simulation.

**S2 Appendix.II Divergence of the lifespan at one side**. Explain how the lifespan predicted by the model switch from finite to infinite at the edge of high ethanol concentration.

**S3 Appendix.III Analytical explanation of the dependence of range** *w* **on parameters**. Analytically explains the dependence of range *w* over feeding amplitude and frequency.

## Acknowledgments

We thank Frank Jülicher and Jochen Guck for helpful discussions. S.P. was supported by funds from TU Dresden’s Institutional Strategy, financed by the Excellence Initiative of the Federal (Saxony) and State (Germany) Governments. T.V.K. acknowledges financial support from the Max Planck Society and X.Z., V.Z., and T.V.K—from Volkswagen Foundation “Life?” initiative.

